# Pulsed Focal ultrasound as a non-invasive method to deliver exosomes in the brain/stroke

**DOI:** 10.1101/2021.02.03.429621

**Authors:** Ahmet Alptekin, Mohammad B Khan, Roxan Ara, Mohammad H Rashid, Fengchong Kong, Mahrima Parvin, Joseph A Frank, Rajiv Chopra, Krishnan Dhandapani, Ali S. Arbab

**Author notes:** **Corresponding author:** Ali S. Arbab, MD, PhD, Georgia Cancer Center, Augusta University, 1410 Laney Walker Blvd, Room CN3315, Augusta, GA 30912, USA.

## Abstract

Exosomes, a component of extracellular vesicles are shown to carry important small RNAs, mRNAs, protein, and bioactive lipid from parent cells and are found in most biological fluids. Investigators have demonstrated the importance of mesenchymal stem cells (MSCs) derived exosomes in repairing stroke lesions. However, exosomes from endothelial progenitor cells (EPCs) have not been tested in any stroke model nor has there been an evaluation of whether these exosomes target/home to areas of pathology. Targeted delivery of IV administered exosomes has been a great challenge and a targeted delivery system is lacking to ***deliver naïve (unmodified) exosomes*** from EPCs to the site of interest. Pulsed focused ultrasound (pFUS) is being used for therapeutic and experimental purposes. There has not been any report showing the use of pulsed low-intensity pFUS to deliver exosomes to the site of interest in models of stroke. In this proof of principle study, we have shown different parameters of pFUS to deliver exosomes in the intact and stroke brain with or without IV administration of nanobubbles. The study results showed that administration of nanobubbles is detrimental to the brain structures (micro bleeding and white matter destruction) at peak negative pressure (PNP) of >0.25 MPa, despite enhanced delivery of IV administered exosomes. However, without nanobubbles, pFUS PNP = 1 to 2 MPa enhances the delivery of exosomes in the stroke area without altering the brain structures.

## Introduction

Exosomes are 30–150 nm lipid bi-layered extracellular bioactive vesicles of endosomal origin that are secreted by all the cells and present in various body fluids. The exosomal content is released by the fusion of these endosomes with the plasma membrane [1]. Exosomes contain lipids, proteins, DNA, RNA, and various metabolites from cells of origin [2]. Their stability in the extracellular environment, ability to carry a payload and specificity to tissues and organs got attention to the use of exosomes as a vehicle to deliver drugs and other treatments to target sites in the body, including the brain [3]. Exosomes have been used for therapeutic purposes to target stroke and other disorders [4, 5]. Investigators have shown the importance of mesenchymal stem cells (MSCs) derived exosomes in the repair of stroke lesions [6]. Repeated administration of exosomes in the rat subjected to the “mechanical occluded” stroke model showed improved functionality and reduced stroke injury [6, 7]. Targeted delivery of IV administered exosomes has been a great challenge, especially through blood-brain-barrier (BBB). Modification of exosomal surface to carry different ligands or peptides have been tried to increase delivery to target tissues [8, 9]. However, the overall results were not encouraging [10, 11]. Investigators are trying to deliver naïve exosomes without surface modification for optimal effect [12]. Delivery of exosomes to the brain is a daunting task and BBB penetrable peptides have been considered [13, 14]. Targeted delivery systems are lacking to ***deliver naïve (unmodified) exosomes*** to the site of interest, especially in the brain or in the area of stroke.

The BBB consists of endothelial and neuronal cells, and it prevents most of the content of the blood from reaching the brain, including most of the drugs and exosomes. Different osmotic and chemical techniques have been developed to overcome BBB with various success, as well as physical disruption by ultrasound. Therapeutic ultrasound delivers mechanical forces that when coupled with ultrasound contrast agent microbubbles can disrupt the BBB and enhances permeability [15]. Moreover, ultrasound contrast agents such as micro/nanobubbles intensify this acoustic force in the vessels and amplify the effect of ultrasound. Ultrasound alone or in combination with micro/nanobubbles have been used to open BBB [16-19] and facilitated the delivery of drugs [20, 21], immunoglobulins [22], albumin [23], and antibodies [24] to the brain. The purpose of the study was to test and optimize the parameters of pFUS with or without nanobubbles to transiently disrupt BBB to enhance the accumulation of IV administered exosomes in the area of interest without damaging the brain parenchyma. The relatively optimized pFUS were then applied to deliver exosomes to lesions in a mouse with transient middle cerebral artery occlusion (MCAO) mediated stroke model.

## Materials and Methods

### Ethics statement

All the experiments were performed according to the National Institutes of Health (NIH) guidelines and regulations. The Institutional Animal Care and Use Committee (IACUC) of Augusta University (protocol #2014–0625) approved all the experimental procedures. All animals were kept under regular barrier conditions at room temperature with exposure to light for 12 hours and dark for 12 hours. Food and water were offered ad libitum. All efforts were made to ameliorate the suffering of animals. CO2 with a secondary method was used to euthanize animals for tissue collection. A minimum of 4 animals was included in each experimental group.

### Chemicals and antibodies

All chemicals and antibodies used in the studies were bought from reputed commercial vendors. Anti-albumin (A0353, ABClonal), and FITC-conjugated AffiniPure Goat Anti-Rabbit (111-095-003, Jackson Laboratories) were used to determine the leakage of BBB. Luxol Fast Blue (#212170250, Acros Organics) used to assess the integrity of white matters, Prussian Blue Stain kit (#3160, Eng Scientific) was used to detect micro bleeding in the brain. HEK293 cells were acquired from ATCC and mouse endothelial progenitor cells (EPC) were collected from the bone marrow of mice using a magnetic sorter as per our previous methods [25]. CellTracker™ CM-DiI (C7000, Thermo Fisher) lipophilic fluorescent dye was purchased for exosome labeling. Fluorescent (FITC) tagged tomato lectin was used to outline the blood vessels (DL-1174, Vector Labs).

### Preclinical bench-top focused UltraSound System (RK-50)

The RK-50 (FUS Instruments Inc, Canada) is a standalone preclinical focused ultrasound system designed for large throughput blood-brain barrier experiments. Atlas-based targeting combined with an optional multi-modality high field insert allows the system to be used with or without magnetic resonance imaging (MRI) or X-ray computed tomography (CT) guidance and imaging. The system is based upon a flexible cross-platform animal handling system simplifying handling and increasing efficiency. The instrument has a three-axis motorized positioning system, accurate stereotaxic-guided targeting using rodent brain atlases (mouse & rat), calibrated spherically focused ultrasound transducer (typically 1.47 MHz) with a maximum RF power of 15W, and includes animal restraint and inhalant anesthetic fixtures. It has custom-written software for atlas registration and treatment planning capability to deliver single or multi-point acoustic exposures (continuous or pulsed).

### Nanobubbles and characterization

Nanobubbles as ultra-sonogram contrast agents were purchased from FUS Instruments Inc. (Toronto, Canada) and prepared according to the supplied protocol. In brief, the unmixed vial was placed in a vial mixture and shaken at 4500 rpm for 5 minutes to make activated emulsion (nanobubbles) containing gas. Then the vial was centrifuged upside down at 800 rpm for 5 minutes. Following centrifugation, the solution containing emulsion of nanobubbles was drawn into a 1 ml insulin syringe without disturbing the foamy upper part. Following activation of nanobubbles, the size and zeta potential of the nanobubbles were determined by a nanoparticle tracking analyzer (NTA). The nanobubbles were diluted 1:1 ratio using sterile PBS before injecting into the tail vein of the mice.

### Parameters of pulsed focal ultrasound (pFUS) in normal brain

We reviewed previous studies (**Table 1**) and set up our parameters. All experiments were performed using a 1.47 MHz (frequency) focal (concave) transducer (wavelength of 1.047 mm). Pulsed ultrasound was applied to 5 points in and around the bregma (at 3 mm lateral from the midline and 3.5 mm deep from the surface on both sides of the brain, with 1% duty cycle (burst duration of 10ms, repetition period 1000ms), for 90 to 180 burst. We have used 0.25 to 2 MPa amplitude of pressure (acoustic power) to the tissue with or without nanobubbles to determine the changes in the BBB and the damages to the structures of the brain tissues. Animals received 0.5 to 2 MPa pFUS to one hemisphere of the brain. After completing pFUS without nanobubbles on one hemisphere, nanobubbles were administered by IV, and 0.25 to 1 MPa pFUS given to the contralateral side of the brain.

**Table 1.** Previous ultrasound studies and parameters to open BBB.

| Reference | Organism | Pressure<br>(MPa) | Frequency<br>(MHz) | Burst<br>rate (Hz) | Burst<br>(ms) | Duration<br>(s) |
| --- | --- | --- | --- | --- | --- | --- |
| [18] | Rabbit | 1 | 1.5-1.63 | 1 | 100 | 20 |
| [26] | Mice | 0.3 | 1.5 | 10 | 6.7 | 60 |
| [27] | Rabbit | 0.4 | 0.69 | 1 | 10 | 20 |
| [28] | Primates | 0.3 | 0.5 | 2 | 10 | 120 |
| [21] | Rats | 0.2 | 1 | 10 |  | 60 |
| [29] | Mouse | 0.3 | 1.5 | 10 | 20 | 60 |
| [30] | Rats | 0.24 | 0.558 | 1 | 1 | 120 |
| [31] | Rats | 0.55 | 0.4–1.7 | 1 | 10 | 60 |
| [32] | Mice | 0.3 | 0.5 | 1 | 10 | 120 |
| [33] | Rats | 0.62 | 0.4 | 1 | 10 | 120 |
| [24] | Mice | 0.3 | 0.558 | 1 | 10 | 120 |
| [34] | Mice | 0.44 | 1.5 | 1 | 10 | 120 |
| [35] | Mice | 0.67 | 1.525 | 10 | 20 | 30 |
| [36] | Rats | 0.3 | 0.548 | 0.5-0.6 | 10 | 120 |
| [23] | Rats | 0.3 | 0.590 | 1 | 10 | 120 |

### Preparation of Animals

We have used 6-8 weeks old immunocompetent Balb/c mice (both males and females) as well as C57BL/6 mice with stroke (24hrs after middle cerebral artery occlusion (MCAO)). Following gas anesthesia with aseptic techniques, a midline incision was made on the scalp to expose and register the points of bregma and lambda. With the help of a mouse brain atlas, we have selected a central point at 3 mm lateral to bregma and 3.5 mm deep in the brain tissue and then selected four more points at 2mm apart from the central point (left, right, front, and back). Then desired pFUS was applied with or without IV nanobubbles administration. Following pFUS, the skin wound was sutured, and the animals were allowed to recover. When applicable, DiI labeled exosomes were IV administered before or soon after the completion of pFUS to determine the distribution of exosomes in the normal and pFUS irradiated areas.

### DiI Labeling of Exosomes

12×10^9^ HEK293 cell-derived exosomes in 1ml PBS incubated with 5µl DiI fluorescent dye at 37°C for 10 minutes followed by incubation at 4°C for 20 minutes. Labeled exosomes washed by adding 1 ml PBS and centrifugation with 30k centrifugal filter (UFC4 LTK 25, Amicon) for 30 minutes at 3900 RPM. After a second wash, exosomes were suspended in 0.6 ml volume of PBS.

### Immunohistochemistry

Luxol fast blue staining was done to determine the integrity of white matter. Staining for extravasated albumin was done to determine the leakage of BBB. Staining for blood vessels was done by FITC-tagged tomato lectin. These stainings were done according to the manufacturer’s protocol. Micro bleeding in the brain was determined by DAB enhanced Prussian blue staining as per our published protocols [37]. To evaluate the tissue integrity, hematoxylin and eosin staining were used.

### Magnetic resonance imaging (MRI)

A 7T dedicated small animal MRI system was used to acquire all the MR images (BioSpec 70/20 USR, Bruker). Animals underwent MRI soon after (within 60 minutes) and at 24hrs following pFUS experiments. To determine the edema formation, T2-weighted, as well as multislice-multi-echo (MSME) T2 images were acquired, followed by the creation of T2-maps. To determine BBB leakage, animals received IP injection of Gd-DTPA contrast agent, and T1-weighted images were acquired. Following the last MRI at 24 hrs, animals were euthanized, perfused, and the brains were collected and fixed for further histochemical studies.

### Making of stroke model, MRI, and pFUS in the stroke areas

Briefly, anesthetized mice have undergone a midline incision on the ventral side of the neck, the right common carotid artery (CCA), the right external carotid artery (ECA), and the right internal carotid artery (ICA) were assessed [38]. Based on the body weight, a silicone rubber-coated monofilament suture (Doccol Corp) was introduced into the ECA lumen and advanced into the ICA until a slight resistance is felt to occlude the origin of the middle cerebral artery (MCA). At 60-min post-occlusion, the filament was gently withdrawn to reinstate cerebral blood flow (CBF) in large vessels as determined with laser speckle imager as described previously [39]. All animals underwent MRI 24hrs after stroke to determine the region and extent of the stroke areas. Based on the MR images, 10 points from trans axial sections of the stroke area were selected for pFUS (from 0.5 to 2 MPa) to facilitate the delivery of exosomes in randomly chosen animals. 90 burst of pFUS applied sequentially to 10 points and 3×10^9^ HEK293 exosome IV administered. Three hours following IV injection of DiI labeled exosomes, all stroke animals were euthanized, perfused with PBS, and the brains harvested to determine the accumulation of exosomes.

### Statistical analysis

Quantitative data were expressed as mean ± standard deviation unless otherwise stated, and statistical differences between more than two groups were determined by analysis of variance (ANOVA) followed by multiple comparisons using Tukey’s multiple comparisons test. Comparison between 2 samples was performed by Student t-test. GraphPad Prism version 8.2.1 for Windows (GraphPad Software, Inc., San Diego, CA) was used to perform the statistical analysis. Differences with p-values less than 0.05 were considered significant(*p<.05,**p<.01, ***p<.001, ****p<.0001).

## Results

### BBB leakage observed on MRI

Characteristics of nanobubbles are shown in **Figure 1**. The size of the nanobubbles was 173.6 (±85.4) nm with a zeta potential of −29.10 ±2.10mV. To determine the effect of applied pFUS PNP and BBB leakage with or without nanobubbles, animals underwent post-contrast T1-weighted MRI 1 hour and 24 hrs after pFUS. Using a stereotactic-guided focused ultrasound device, we tested different pFUS PNP to find the maximum non-disruptive dose. We applied pFUS into the right hemisphere without nanobubbles and the left hemisphere with nanobubbles as indicated above. pFUS in the right hemisphere without nanobubbles was applied first, then pFUS was applied on the left hemisphere during infusion of nanobubbles. pFUS parameters were consisted of 180 cycles, with 10 ms burst and 1000 ms pulse repetition time of each cycle. The pFUS dose ranges from 0.5 to 2 MPa without nanobubbles did not cause any BBB leakage indicated by non-enhancement at the site of pFUS in the right hemisphere (**Figure 2 A-C**). On the other hand, 1-hr MRI images showed BBB leakage at the site of pFUS with nanobubbles even at the lowest dose of 0.25 Mpa, although the leakage at 0.25 MPa was transient as no enhancement was observed on 24-hr postcontrast MRI (**Figure 2A**). pFUS doses with nanobubbles at 0.5 and 1 MPa showed sustained BBB leakage, which was observed on 24-hr MRI and contrast-enhanced areas on the left hemisphere (**Figure 2 B-C**, left hemispheres). A significant advantage of using nanobubbles was observed for opening BBB, even with the lowest dose applied. We also observed a decrease in the signal intensity following contrast administration on MRI at 24hrs in animals that received 0.5 MPa and 1 MPa pFUS with nanobubbles (**Figure 2B, C**), which might indicate a gradual repair of BBB leakage.

**Figure 1.**
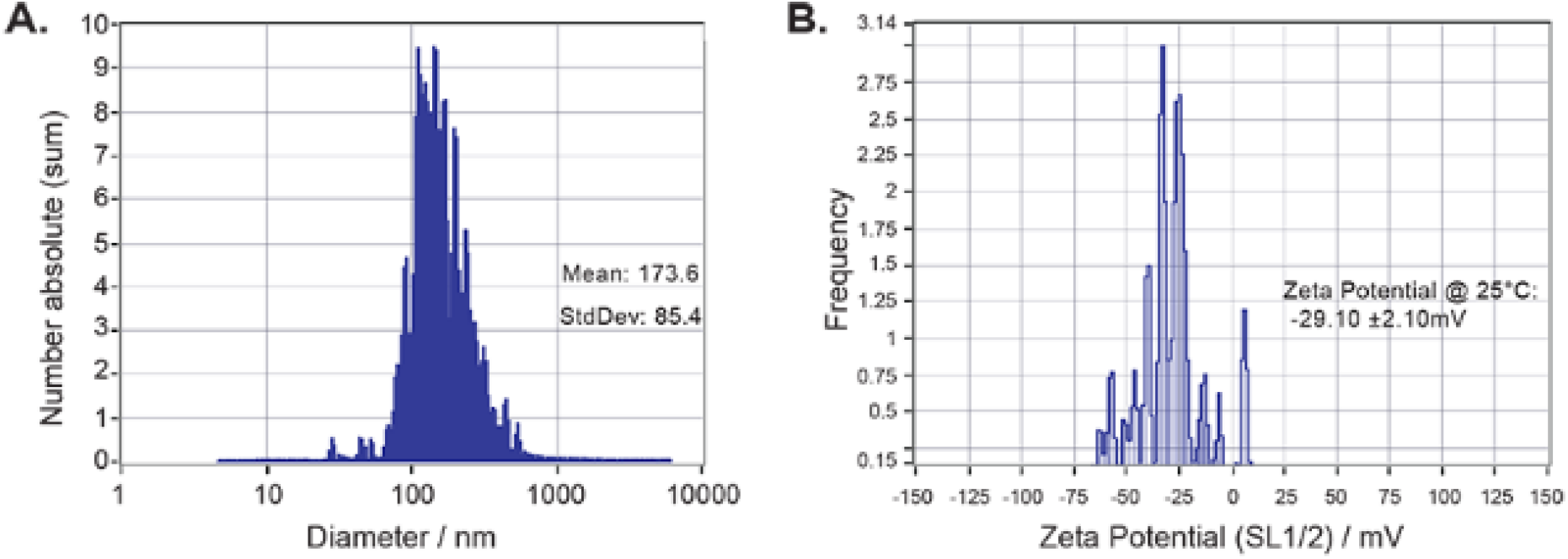
NTA analysis of nanobubble’s size (**A**) and zeta potential (**B**).

**Figure 2.**
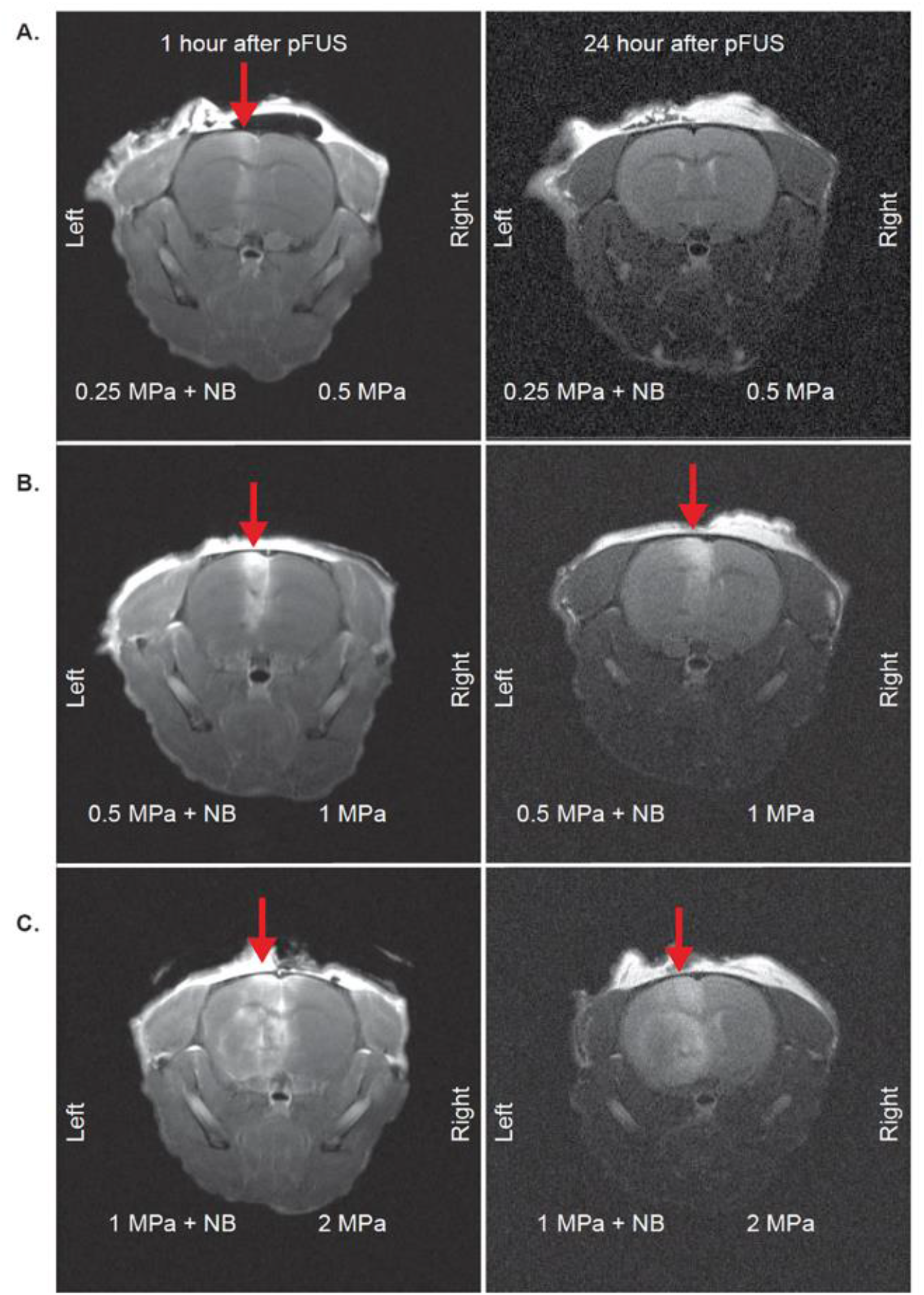
Post-contrast T1 images to evaluate BBB opening of mice brains with pFUS application. 1 hour and 24 hour MRI images of mice with 0.25MPa + NB to left hemisphere, 0.5MPa to right hemisphere (**A**); 0.5MPa + NB to left hemisphere, 1MPa to the right hemisphere (**B**); 1MPa + NB to left hemisphere, 2MPa to the right hemisphere (**C**). Red arrows indicate leakage from BBB.

### Damage of intact brain following pFUS

To evaluate tissue damage and bleeding after pFUS application, we sacrificed all animals after 24 hours of pFUS, and harvested brain tissue, fixed it with paraformaldehyde, and made paraffin blocks to make sections for different IHC analysis. We stained tissues with luxol fast blue (LFB) and Prussian blue to evaluate the damage to white matter (myelin sheath) and bleeding, respectively. LFB staining showed damage to the myelin sheath or white matter, starting with 0.5 MPa pFUS when nanobubble was used (**Figure 3B-C**, left). However, we did not observe similar damage to the white matter even with 2 MPa pFUS when nanobubbles were not used (**Figure 3 A-C**, right). Similar to the damage of white matters, micro bleeding (DAB enhanced Prussian blue staining, brown spots) was also observed in the pFUS irradiated area (both at 0.5 and 1 MPa) when nanobubbles were used (**Figure 4 B-C**, left). On the other hand, no micro bleeding was observed even with the highest pFUS (at 2 MPa) without nanobubbles (**Figure 4 A-C**, right). These findings, in combination with MRI findings, indicate that 0.25 MPa with nanobubbles opens BBB without causing structural damage or bleeding, but using pFUS PNP > 0.5 MPa with nanobubbles caused structural damage to the intact brain with the parameters described above. However, pFUS without nanobubbles is safe even with the highest applied 2 MPa intensity.

**Figure 3:**
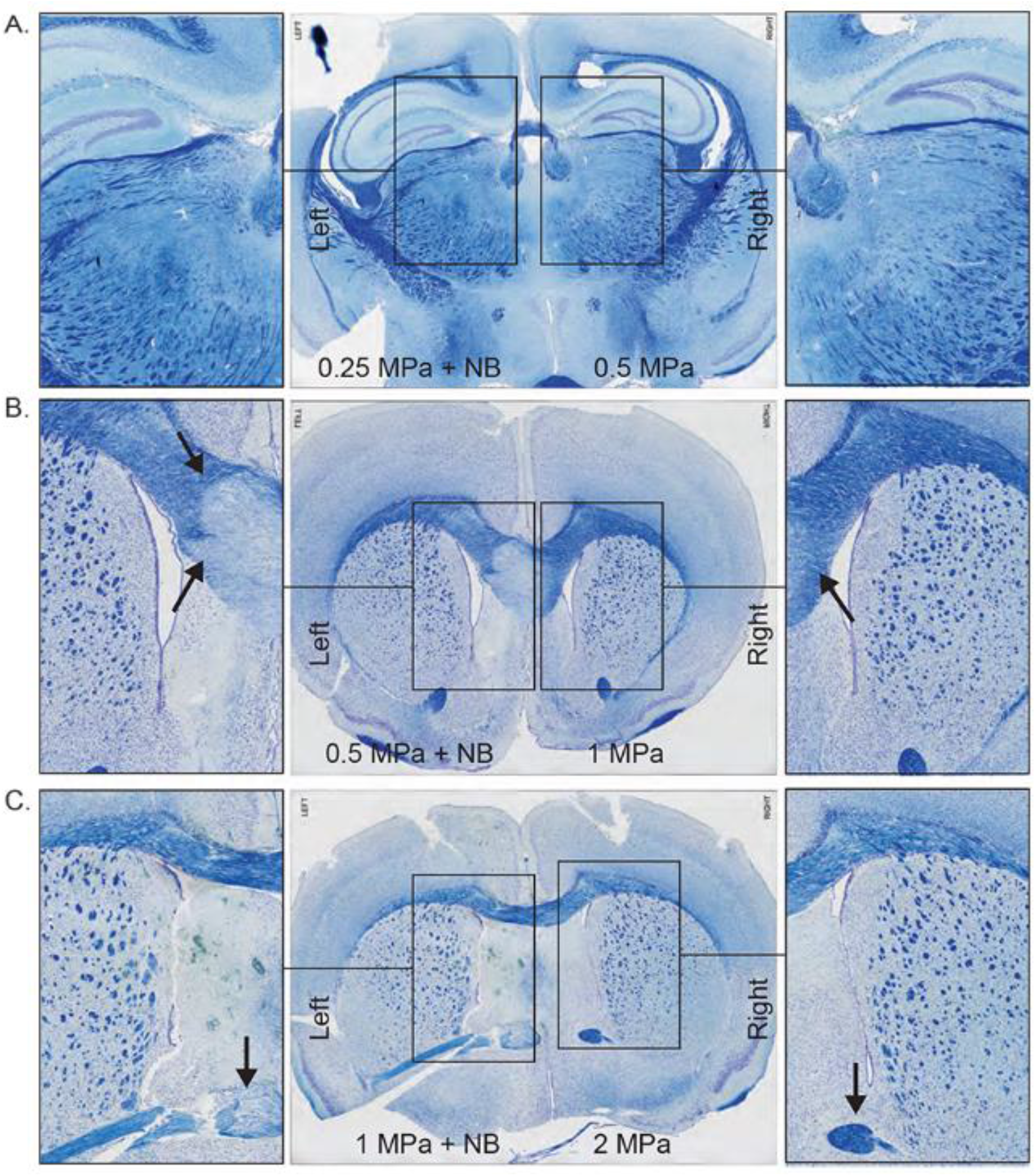
Luxol Fast Blue staining of pFUS applied mice brain with 0.25MPa + NB to left hemisphere, 0.5MPa to the right hemisphere (**A**); 0.5MPa + NB to left hemisphere, 1MPa to the right hemisphere (**B**); 1MPa + NB to left hemisphere, 2MPa to the right hemisphere (**C**).

**Figure 4:**
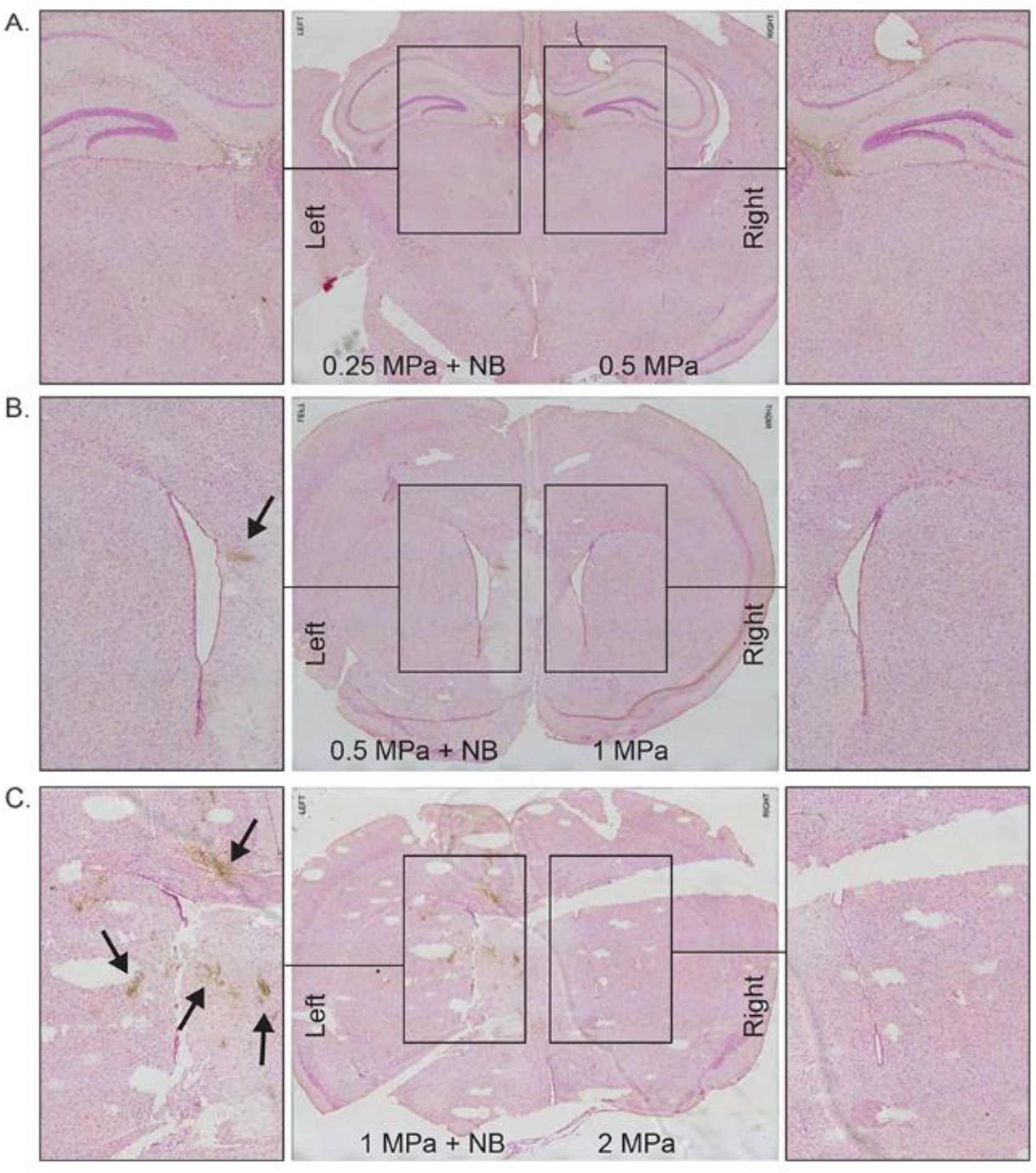
Prussian Blue staining of pFUS applied mice brain with 0.25MPa + NB to left hemisphere, 0.5MPa to the right hemisphere (**A**); 0.5MPa + NB to left hemisphere, 1MPa to the right hemisphere (**B**); 1MPa + NB to left hemisphere, 2MPa to the right hemisphere (**C**). Arrows indicate hemosiderin stained with Prussian blue.

### Confirmation of BBB opening

Albumin is a plasma protein and its leakage into brain parenchyma indicates a BBB opening [40]. To investigate albumin leakage, we stained the paraffin sections with FITC-anti-albumin antibody. Leakage of albumin (labeling of FITC-tagged anti-albumin antibody) was observed when pFUS was used following IV administration of nanobubbles only in cases where pFUS intensity was at 1 MPa and 0.5 MPa (**Figure 5 A-C**, left). However, we have not observed any leakage of albumin following irradiation up to 2 MPa pFUS without nanobubbles (**Figure 5A-C**, right). Corresponding HE staining showed no obvious damage/necrosis to the area of albumin leakage (**Figure S1**).

**Figure 5:**
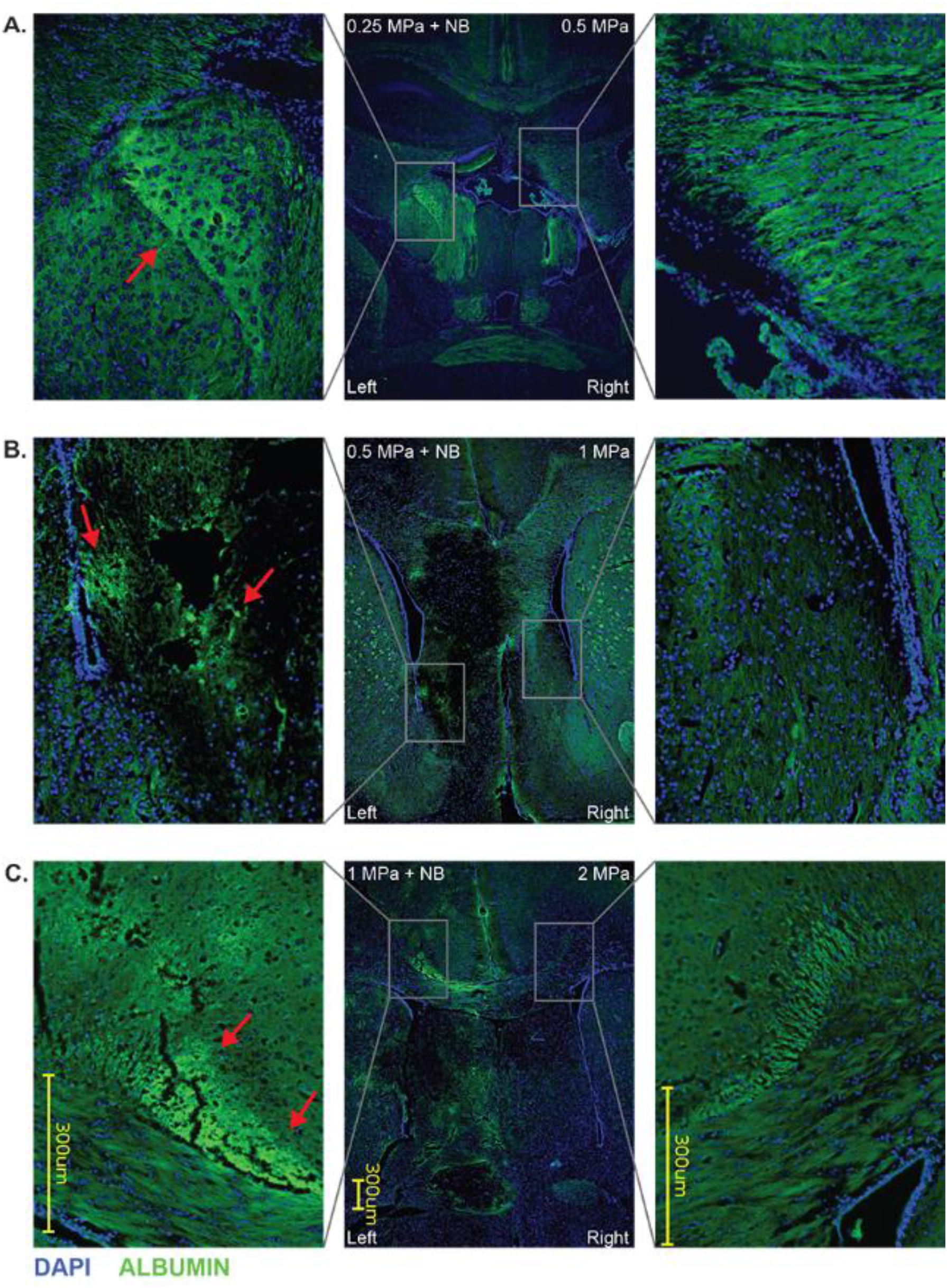
Albumin and DAPI immunofluorescence labeling of pFUS applied mice brain with 0.25MPa + NB to left hemisphere, 0.5MPa to the right hemisphere (**A**); 0.5MPa + NB to left hemisphere, 1MPa to the right hemisphere (**B**); 1MPa + NB to left hemisphere, 2MPa to the right hemisphere (**C**). Arrows indicate fluorescent-labeled albumin.

### Exosome delivery into the brain in the stroke model

In a cerebral stroke, the BBB is already compromised. Our previous studies showed the accumulation of neural stem cells (NSC) derived exosomes in the stroke areas without the application of pFUS [41]. We wanted to see whether the application of pFUS without nanobubbles will enhance the accumulation of IV administered exosomes in the stroke areas. We used the right middle cerebral artery occlusion (MCAO) as a stroke model in mice. Twenty-four hours after stroke, we obtained MRI (**Figure 6 A-C**, middle). After determining the stroke area, we applied 0.5 MPa (**Figure 6A**), 1 MPa (**Figure 6B)**, and 2 MPa (**Figure 6C**) pFUS to 10 selected points in the stroke area to induce BBB opening and administered DiI-labeled HEK293 derived exosomes via IV injection. Because the stroke already resulted in damage to the parenchyma, further disruption with nanobubbles addition would be detrimental and therefore nanobubbles were not infused. A second IV dose of exosomes was administered the next day following pFUS alone, and brains were harvested 3-hours post-sonication. We found the accumulation of exosomes was enhanced when pFUS was applied at the area around the stroke sites (**Figure 6A-C;** right), compared to corresponding sections in the contralateral side of the brain (**Figure 6A-C;** left). We also observed significantly (p<0.01) increased exosome delivery in animals that received pFUS at PNP of 1MPa and 2MPa pFUS application, compared to 0.5MPa. The number of exosomes in respect of the total area of pixels was determined by the color threshold program using ImageJ. 2 MPa PNP showed the total area of exosomes (0.12±0.03) vs. (0.02±0.03) for 0.5 MPa. 1 MPa showed (0.10±0.10) area of pixels. These findings indicate that pFUS without nanobubbles could be used as a method to induce the delivery of exosomes into brain parenchyma.

**Figure 6:**
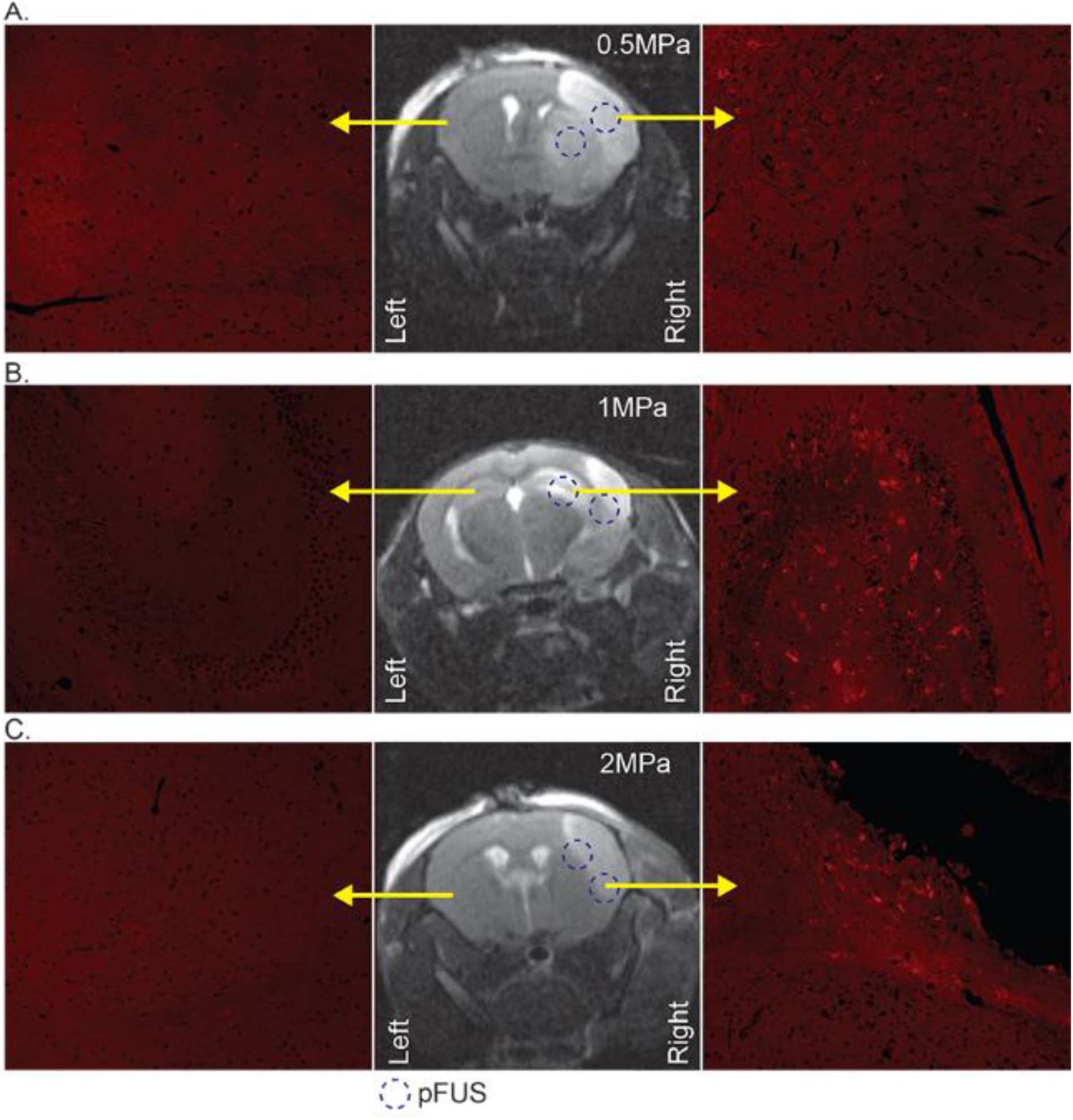
MRI images (middle column) to indicate stroke area of mice and dotted circle to delineate the 2 of pFUS application points. The right column with fluorescent microscopy images to show DiI-labeled (red) exosomes from the right hemisphere of the mice with 0.5MPa (**A**), 1MPa (**B**), and 2MPa (**C**) pFUS application; left columns with fluorescent microscopy images to show the contralateral side of the brain without stroke and pFUS application.

We checked whether the exosomes are attached to the endothelial lining or extravasated and accumulated in the brain parenchyma. Sections were stained with FITC tagged tomato lectin to delineate the endothelial lining in the area of exosome accumulation. Although the exosomes were found inside the vessel, high-resolution images showed the accumulation of exosomes in the deeper part of the brain parenchyma away from the endothelial lining. This finding indicates blood vessels’ opening to allow the exosomes’ accumulation into the brain (**Figure 7**).

**Figure 7:**
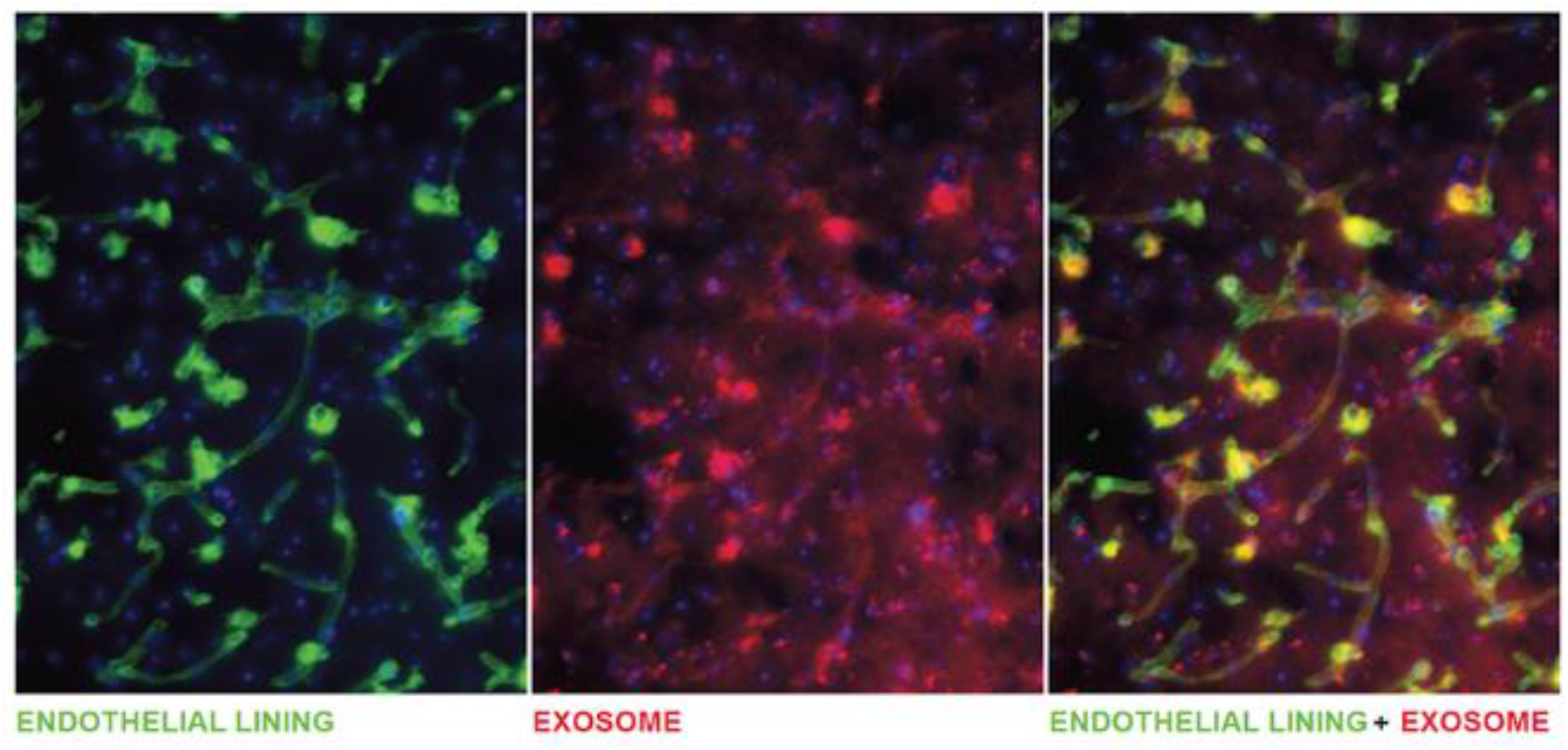
Fluorescent image of FITC-labeled Lectin delineates endothelial lining of vessels and DiI positive exosomes.

## Discussion and notes

The study’s purpose was to determine the safe pFUS parameters that will allow delivering the naïve exosomes at the stroke site. Our studies clearly showed that pFUS at PNP of 1 to 2 MPa could enhance the delivery of naïve intravenously administered exosomes in the stroke areas. Many studies have utilized pFUS to deliver nanoparticles or other agents at the sites of irradiation (pFUS) with or without administration of microbubbles [42]. Previous studies showed enhanced delivery of nanoparticles into intact brain parenchyma when pFUS was used [43]. However, none has used the technique to deliver exosomes to the site of a stroke. It is already known that BBB is already open at the site of stroke [44], which may preclude the use of nano/microbubbles during pFUS. The use of nano/microbubbles may be detrimental to the stroke area. Moreover, stroke area showed extensive inflammatory process due to accumulation of neutrophils and other inflammatory cells [45], therefore causing more inflammation may not be an expected outcome following pFUS mediated injury [23].

The current stroke treatment is thrombolysis in the stroke’s acute phase [46]. However, more treatment options and safe delivery of therapeutics into brain parenchyma are needed. One strategy is to deliver stem cell-derived extracellular vesicles to facilitate the repair or stimulate the replacement of the infarcted tissue. It has been shown that there was no difference in recovery after ischemic stroke in mice following the infusion of mesenchymal stem cells (MSC) as compared to extracellular vesicles derived from the MSC [47]. Another study demonstrated extracellular vesicles from MSC also improved recovery of stroke in rat models [48]. Previously, we have reported that extracellular vesicles from human neural stem cells have improved the recovery in a stroke model in mice [41]. Taken together, these studies demonstrate that exosomes could be a novel therapeutic in the treatment of stroke. However, all of the studies mentioned above relied on the passive distribution and accumulation of exosomes at the penumbra or peri-stroke regions.

While the BBB is responsible for excluding large molecules, it can allow the transport of small lipophilic molecules (< 400-500 Da) [49]. This makes the brain out of reach for most drugs, including most of the exosomes. Among different strategies to overcome BBB to reach the brain, it has been demonstrated that pFUS mechanical forces can disrupt and open BBB without permanent damage to brain parenchyma in rodents [19]. By optimizing this technique with microbubbles and employing MRI guidance, researchers successfully opened the BBB in patients with Alzheimer’s [50] and amyotrophic lateral sclerosis [16]. It has also been demonstrated that magnetic resonance-guided pFUS with MB resulted in BBB opening in glioma patients and increased delivery of chemotherapeutics [51]. To date, radiotherapy and chemotherapy agents, antibodies, gene therapy agents, nanoparticles, and cells were able to be transferred to the brain with the help of pFUS coupled with MB infusion [52].

In the treatment of stroke, therapeutic ultrasound (sonothrombolysis) has been tested as an adjunctive to tPA therapy. Although it was found to be safe, there was no clinical benefit in stroke patients [53]. There is presently an ongoing phase 2 clinical study that is investigating the use of therapeutic ultrasound in combination with microbubble infusion (NCT01678495). Besides these thrombolytic approaches in stroke, other studies have employed therapeutic ultrasound to deliver therapeutics to brain parenchyma. Previous studies showed that enhanced delivery of nanoparticles in the brain parenchyma and in the tumor areas when pFUS was applied without any microbubbles. This enhanced delivery was due to the lowering of interstitial pressure [43, 54].

The transient nature of BBB opening was also observed following low-intensity pFUS+MB, and it has been reported that the BBB closes at 28h [35], 24h [29], 8 [26]. This indicates that pFUS does not cause permanent damage and can temporarily open BBB for therapeutic delivery in the intact brain. Clinical trials demonstrated ultrasound-induced BBB opening was closed after 24h and proved its safety in humans to be used in therapeutic delivery into the brain [50]. Based on our findings, pFUS at 0.25 MPa can be used to open BBB and deliver exosomes without damaging the brain structures.

However, our aim was to increase the delivery of exosomes in the stroke area without nanobubbles administration. It is already known that BBB is already open in a stroke area [44]. Therefore, we tried to find the best possible parameters to deliver naïve IV administered exosomes to the stroke site without destructing the brain structures. As our results showed no detrimental effects in the intact brain, we have used pFUS with 0.5 to 2 MPa intensities without nanobubbles. Our results indicated any intensities above 1 MPa would allow enhanced delivery of naïve IV administered exosomes in the stroke areas away from the endothelial lining. We used HET293 derive exosomes, and we expect that exosomes derived from EPCs would accumulate more in the area of stroke following low pFUS. We are now investigating the delivery of EPC derived exosomes and their effect on stroke outcomes.

## Limitation of the studies

We have not studied the expression of different cytokines following different doses of pFUS. Previous studies showed sterile inflammatory response following pFUS in the brain parenchyma [23]. However, it is already known that stroke caused a massive inflammatory reaction. Our future studies will be measuring inflammatory cytokines using membrane array or proteomics in the stroke areas with or without pFUS. It is also known that higher PNP can cause heating of the skull at the irradiation site [55]. Moreover, local heating may cause exosome accumulation under the skull rather than inside the brain parenchyma. However, our IHC analysis showed an accumulation of exosomes in the deeper part of the brain parenchyma. Ultrasound may alter the neurobehaviour of the animals [56]. Our next sets of studies are looking at the neurobehaviour of animals following pFUS with or without EPC exosomes. The results are forthcoming.

## Conclusion

pFUS without micro/nanobubbles can be used to deliver IV administered exosomes to the site of interest. Engineered or non-engineered exosomes can be used as therapeutics for stroke therapy in addition to tPA.

## Acknowledgment

The study was funded by American Heart Associate (AHA) grant 19TPA34850076 and part of Georgia Cancer Center start-up fund to ASA, and NIH grant R01NS117565 to KD.

