## Supplemental Figure 1 for "Pulsed Focal ultrasound as a non-invasive method to deliver exosomes in the brain/stroke"

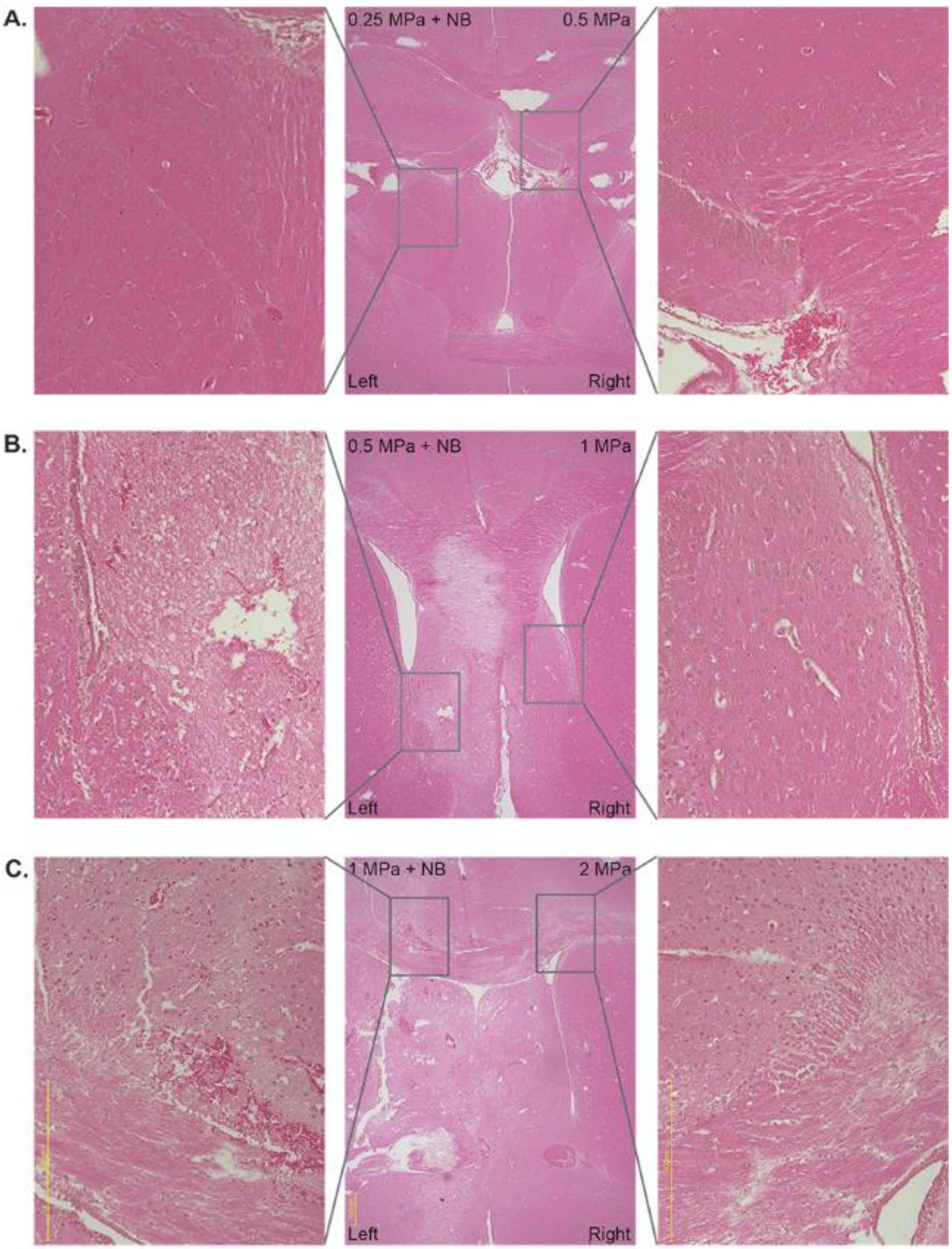


**Figure S1:** Hematoxylin and eosin staining of pFUS applied mice brain with 0.25MPa + NB to left hemisphere, 0.5MPa to the right hemisphere (**A**); 0.5MPa + NB to left hemisphere, 1MPa to the right hemisphere (**B**); 1MPa + NB to left hemisphere, 2MPa to the right hemisphere (**C**).
